# Under pressure: integrated endothelial cell response to hydrostatic and shear stresses

**DOI:** 10.1101/2024.05.30.596749

**Authors:** Christian J. Mandrycky, Takashi Ishida, Samuel G. Rayner, Adam Heck, Brandon Hadland, Ying Zheng

**Affiliations:** Department of Bioengineering, University of Washington, Seattle, WA 98195; Institute for Stem Cell and Regenerative Medicine, University of Washington, Seattle, WA 98109; Translational Science & Therapeutics Division, Fred Hutchinson Cancer Center, Seattle, WA 98109; Center for Lung Biology, Department of Medicine, University of Washington, Seattle, WA 98195; Division of Pediatric Hematology/Oncology, University of Washington School of Medicine, Seattle, WA 98195

**Keywords:** hemodynamic pressure, endothelial cells, flow response, arterial fate

## Abstract

Blood flow within the vasculature is a critical determinant of endothelial cell (EC) identity and functionality, yet the intricate interplay of various hemodynamic forces and their collective impact on endothelial and vascular responses are not fully understood. Specifically, the role of hydrostatic pressure in the EC flow response is understudied, despite its known significance in vascular development and disease. To address this gap, we developed in vitro models to investigate how pressure influences EC responses to flow. Our study demonstrates that elevated pressure conditions significantly modify shear-induced flow alignment and increase endothelial cell density. Bulk and single-cell RNA sequencing analyses revealed that, while shear stress remains the primary driver of flow-induced transcriptional changes, pressure modulates shear-induced signaling in a dose-dependent manner. These pressure-responsive transcriptional signatures identified in human ECs were conserved during the onset of circulation in early mouse embryonic vascular development, where pressure was notably associated with transcriptional programs essential to arterial and hemogenic EC fates. Our findings suggest that pressure plays a synergistic role with shear stress on ECs and emphasizes the need for an integrative approach to endothelial cell mechanotransduction, one that encompasses the effects induced by pressure alongside other hemodynamic forces.

## Introduction

Blood flow is a critical driver for the development of functional vasculature to support the growth and homeostasis of tissue and organs during both early development and adulthood. Endothelial cells (ECs) form the innermost layer of blood vessels and are exposed to numerous forces imposed by flowing blood including shear stress, mechanical stretch, and hydrostatic pressure, all of which are fundamental in determining EC fate, modulating EC signaling pathways, and guiding vascular network formation.^1–6^ In the early stages of vascular development, following the initiation of the heartbeat, specific endothelial populations lining the dorsal aorta are subject to shear stress and other mechanical forces, promoting an endothelial-to-hematopoietic transition to establish hematopoietic stem cells responsible for life-long blood production.^7–9^ This transition is intricately regulated by various signaling pathways, such as Notch and KLF2/NO, which are influenced by fluid flow dynamics and play pivotal roles in arterial specification and hematopoietic development.^10–13^ As the heart develops and blood volume increases, ECs orchestrate remodeling of the vascular plexus and the transition to vascular homeostasis.^14–16^ Consequently, ECs transition from the “activated” state seen during development to a quiescent state with reduced proliferation, forming a vascular barrier to regulate oxygen and nutrient exchange and support organ function.^17–19^ Proper flow is also essential for maintaining key signaling pathways in the established vascular endothelium.^20,21^ In adult arteries, flow changes or disturbance may activate NADPH oxidase and promote ROS generation to enhance Notch signaling, influencing angiogenesis and vascular remodeling.^22,23^

Among hemodynamic forces, shear stress has been extensively studied in EC mechanobiology,^14^ regulating cellular alignment,^24,25^ morphogenesis,^26^ angiogenesis,^27^ and promoting an arterial phenotype ^13^ as well as hemogenic fate.^8,9^ Exposure of iPSC-derived ECs to shear stress induces arterial specification, highlighting the critical role of biomechanical forces in endothelial cell fate determination.^28–30^ Shear stress also regulates gene expression in ECs, influencing vascular identity, with laminar flow upregulating atheroprotective genes and disturbed flow increasing atheroprone genes.^31,32^ In contrast, the impact of hydrostatic pressure on EC fate and function has not been as well studied. Elevated hydrostatic pressure affects factors like cell stiffness and proliferation,^23,33,34^ and plays a direct role in the pathogenesis of hypertensive disorders of both the systemic and pulmonary circulation.^35–37^ Hydrostatic pressure gradually increases concomitant with blood flow following the onset of the heartbeat in early embryonic development; however, little is known about the role that pressure may play in mediating embryonic EC fate decisions, and how ECs integrate cues from multiple hemodynamic forces to determine their fates. Further study of the precise effects of each of these hemodynamic forces on EC signaling is critically important for understanding developmental processes and disease mechanisms. In vitro microfluidic models provide an appealing platform for such studies, allowing careful tuning of each flow-related force, both individually and in combination.

In this study, we used a custom microfluidic system to independently control the pressure and shear conditions affecting ECs cultured under flow, using imaging, cell collection, and RNA analysis as readouts to assess EC morphology and transcriptional profiles in response to specific hydrodynamic conditions. Our findings demonstrate that increases in hydrostatic pressure at a constant shear stress, significantly influence EC shape, alignment, and transcriptional response to flow. Translating these findings into a mouse model, we identified that a pressure-responsive gene signature identified in our in vitro microvessel models was conserved in arterial/hemogenic ECs in the developing murine aorta, implicating a role for pressure in EC maturation in vivo. This study provides insights into how ECs integrate mechanical cues, affecting their transcriptome, identity, and function, thereby contributing to the understanding of vascular development and disease progression.

## Results

### Pressure modifies EC morphological responses to flow

To study the influence of pressure on the EC response to flow, we engineered a microfluidic flow circuit comprising a rectangular cross-section culture channel and a resistor channel in series (see Methods), connected to a syringe pump. The design of the flow circuit is depicted in Fig. 1A. By precisely controlling the flow rate from the syringe pump and the geometry of the resistance channel, we achieved individual control of shear stress and pressure within the system. Through brightfield and fluorescence microscopy, we observed distinct morphological and organizational differences in ECs cultured under varying pressure conditions (Fig. 1B, and Supplementary Fig. 1). Under low pressure (Fig. 1B-i), cells were larger and had fewer nuclei per field compared to those under high-pressure conditions. Cells under low pressure showed greater alignment, primarily elongating in the direction of fluid flow. Conversely, under high pressure (Fig. 1B-ii), cells appeared rounder (lower aspect ratio) and demonstrated a more varied orientation, often aligning orthogonally to the flow direction. Significant differences were observed in cell area, alignment angle, and aspect ratio between 0 and 60 mmHg at a shear stress of 5 dyne/cm^2^ (Fig. 1C). However, the percentage of nuclei with detectable Ki67 staining did not significantly differ between the groups (0 mmHg = 62.9 ± 4.5%, 60 mmHg = 63.6 ± 2.9%). In both pressure conditions Notch1-ECD staining polarized towards the downstream cell border while DLL4 staining was diffused across the cell (Fig. 1D).

**Figure 1.**
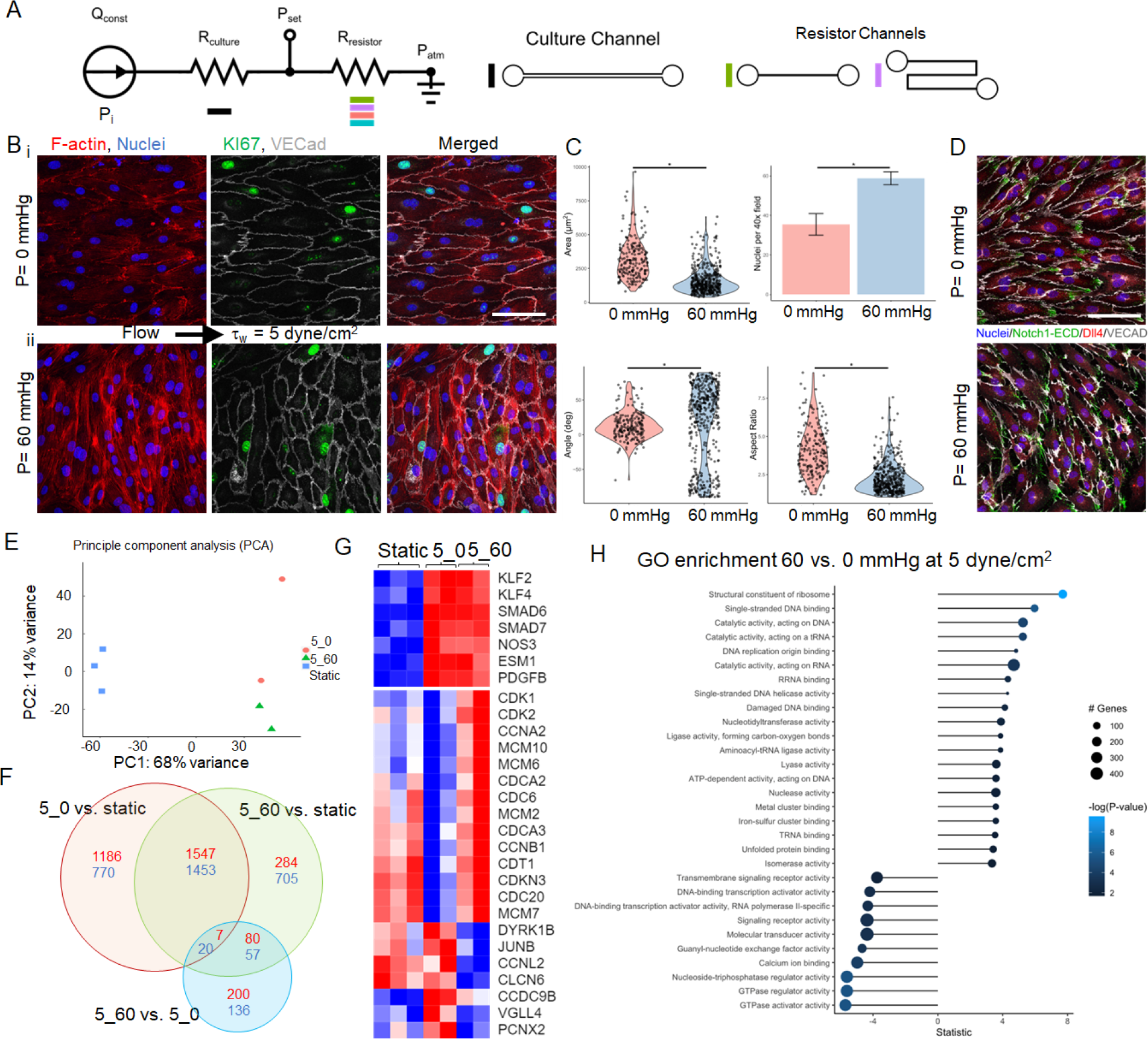
**Microfluidic culture of endothelial cells to decouple pressure and flow effect**. **A**. Schematic of the flow circuit connecting the culture channel and a resistor channel in serial with the syringe pump. Different resistor channels were designed with specific width and height to change the hydrostatic pressure P_set_ at the outlet of culture channels. **B**. Representative images of endothelial cells cultured at shear stress τ_w_ = 5 dyne/cm^2^ show distinct cell morphology and alignment for P_set_ = 0 (i) and 60 mmHg (ii). Red: F-actin; green: KI67; blue: nuclei; gray: VECad. **C**. Quantification of cell area, alignment angle, aspect ratio and number of nulcei per 40x field for to pressure conditions. *: p<0.05. **D**. Representative immunostaining images of endothelial cells shows different polarization of Notch1-ECD (green) at different pressure conditions under flow. **E-H.** Transcriptional changes of ECs in response to flow and pressure. **E.** Principal component analysis showing the separation of static and flow condition in PC1 and mild pressure separation in PC2. **F.** Venn Diagram of differentially expressed gene numbers comparing the effect of flow and pressure on EC response. Red: upregulated genes; blue: downregulated genes. 5_0: τ_w_ = 5 dyne/cm^2^ and P_set_ = 0 mmHg; and 5_60: τ_w_ = 5 dyne/cm^2^ and P_set_ = 60 mmHg. **G.** Heatmap of selected flow-responsive markers (upper) and differentially expressed cell-cycle related genes comparing static and pressure groups. Colormap: log2(CPM) with minimum value in blue and maximum in red. **H**. GO term analysis and the number of genes associated comparing high vs. low pressure at the same flow shear condition.

### Pressure alters the transcriptional profiles of ECs in response to flow

We performed bulk RNA sequencing of ECs cultured in round microfluidic tubes (see Methods) at a flow rate of 5 dyne/cm^2^ and either low (0 mmHg) or high (60 mmHg) pressure conditions, in comparison to cells cultured in wells (∼8 mm diameter) under minimal hydrostatic load (∼0.3 mmHg) without bulk flow for baseline comparison. A cylindrical geometry was chosen to ensure uniform shear and pressure exposure for all collected cells. Principal component analysis (PCA) revealed a clear differentiation between static and flow-exposed samples along PC1, while varying pressure conditions exhibited slight separation along PC2 (Fig. 1E). Under both low and high pressure, flow at 5 dyne/cm^2^ shared overlap of 1559 upregulated and 1473 downregulated genes (3032 total) compared to the static condition (Fig. 1F). Notably, all flow conditions upregulated classical flow-responsive markers (e.g., *KLF2*, *KLF4*, *SMAD6*, *SMAD7*, *NOS3*) regardless of pressure (Fig. 1G), with corresponding gene regulation pathways consistent with established flow responses ^38^ (Supplementary Fig. 2).

Comparison of the two pressure conditions under flow identified 500 differentially expressed genes, the majority of which were not differentially regulated in the transition from the static to perfused conditions (Fig. 1F). Particularly, a subset of genes associated with cell proliferation and cytoskeletal structure were upregulated in the high-pressure condition, most of which remained unchanged or downregulated in response to flow (Fig. 1G, Supplementary Fig. 3). Gene Ontology (GO) analysis revealed upregulation of pathways associated with DNA binding, replication, and activity under high pressure, while pathways related GTPase regulator, activator and calcium ion binding, and molecular transducer activity were downregulated (Fig. 1H, Supplementary Fig. 2). Altogether, these results support the concept that dynamic integration of fluid shear stress and hydrostatic pressure elicits distinct transcriptional responses of ECs, impacting pathways involved in EC proliferation state, signal transduction, and regulation.

### Single-cell RNA-seq reveals a strong transcriptional response to flow, further modified by hydrostatic pressure

To dissect the combinatorial influence of flow and pressure on the EC transcriptome and to probe the heterogeneity in transcriptional responses among individual ECs to these hemodynamic forces, we performed single cell RNA sequencing (scRNA-seq) on over 5,000 ECs subjected to four culture conditions in round channels: static (no flow), or 24 hours of flow shear stress of 5 dyne/cm^2^ at pressures of 0 mmHg, 20 mmHg, and 60 mmHg. Cells from each condition, pooled from multiple devices and individually barcoded, then subjected to parallel processing on the 10X Genomics platform to minimize batch effects. Following the removal of low-quality cells and deconvolution of sample barcodes, dimensionality reduction and clustering using the Monocle3 toolkit revealed three major clusters. This analysis underscored the significant transcriptional impact of shear stress (Fig. 2A). Static samples primarily clustered separately (Cluster 2), while cells exposed to flow shear stress predominantly formed a distinct cluster (Cluster 1) irrespective of pressure. A third cluster (Cluster 3) was characterized by cells expressing genes indicative of active cell cycle (MKI67) (Fig. 2A). The proportion of cells in cluster 3 decreased in response to flow but increased with pressure. Further increments in pressure did not alter the proportion of cells in active proliferative states.

**Figure 2.**
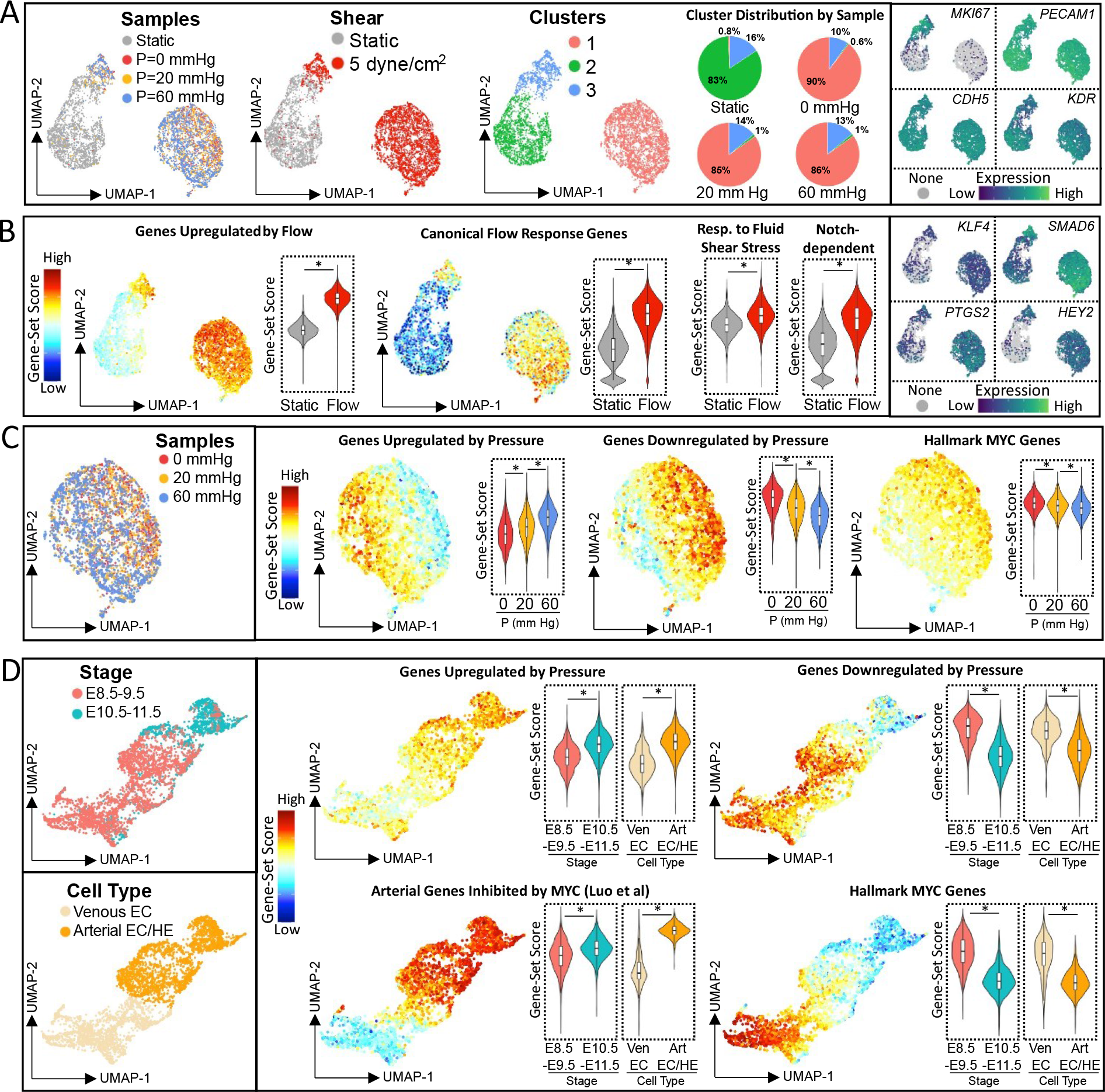
**Single cell transcriptomics analysis of HUVECs subjected to varying flow and pressure parameters reveals a pressure-responsive transcriptional program conserved during murine embryonic vascular development**. **A**. Single cell transcriptomic data from human ECs subjected to static (no flow) conditions or constant flow shear (5 dyne/cm^2^) with varied hydrostatic pressure (P = 0, 20, and 60 mmHg). Shown are UMAP projections with cells classified by Louvain cluster (upper left panel), static vs combined flow conditions (upper right panel), static vs flow conditions separated by pressure (lower left panel), and by expression of pan-endothelial genes (*PECAM1, CDH5, KDR*) or cell cycle gene, *MKI67* (lower right panels) **B**. UMAP projects and violin plots showing aggregated gene-set scores for canonical flow response genes (upper left panel), genes classified in Response to Fluid Shear Stress by Gene Ontology (lower left panel), Notch-dependent genes (upper right panel), and genes experimentally determined as upregulated by flow based on differential gene expression in the current study (bottom right panel). (* p< 2.2E-16, Wilcoxon) **C**. UMAP projection of cells from cluster 1 (flow conditions, MKI67 negative) classified by exposure to pressure (P = 0, 20, or 60 mm Hg) (left panel), and aggregated gene-set scores for genes experimentally determined as upregulated or downregulated by pressure based on differential gene expression in the current study (central panels) and genes classified as Hallmark MYC Target Genes (right panel). (*p< 8E-9, Wilcoxon). **D**. Single cell transcriptomic data from endothelial populations isolated from murine embryos classified by stage (embryonic day, E8.5-E9.5 vs E10.5-E11.5) or by cell type (venous EC vs arterial/hemogenic EC) (left panels, from published data sets^44,45^). UMAP projects and violin plots showing aggregated gene-set scores for genes experimentally determined as upregulated or downregulated by pressure in human ECs in the current study (top panels), arterial genes negatively regulated by MYC (lower panel^39^), and genes classified as Hallmark MYC Target Genes V2 (lower right panel). (*p< 2.2E-16, Wilcoxon).

Uniform expression of pan-endothelial markers, including *CDH5*, *PECAM1*, and *KDR* was consistent across clusters (Fig. 2A). The gene expression disparity between clusters 1 and 2 was driven largely by flow-responsive genes (e.g., *KLF4*, *SMAD6, SMAD7*, *PTGS2*) and Notch-dependent genes (e.g., *HEY1*, *HEY2*, *JAG1*, *SOX17*) (Fig. 2B). Gene-set expression scores for canonical flow/shear stress-response genes (*KLF2*, *KLF4*, *SMAD6*, *SMAD7*, *PTGS2*, *NOS3*) and genes involved in the GO term “Cellular Response to Fluid Shear Stress” were notably elevated under flow conditions (Fig. 2B). Differential gene expression analysis between ECs exposed to flow vs static conditions confirmed the detection of known flow-response genes (Fig. 2B and Table 1), affirming the sensitivity of our scRNA-seq data to hemodynamic stimuli.

**Table 1:**
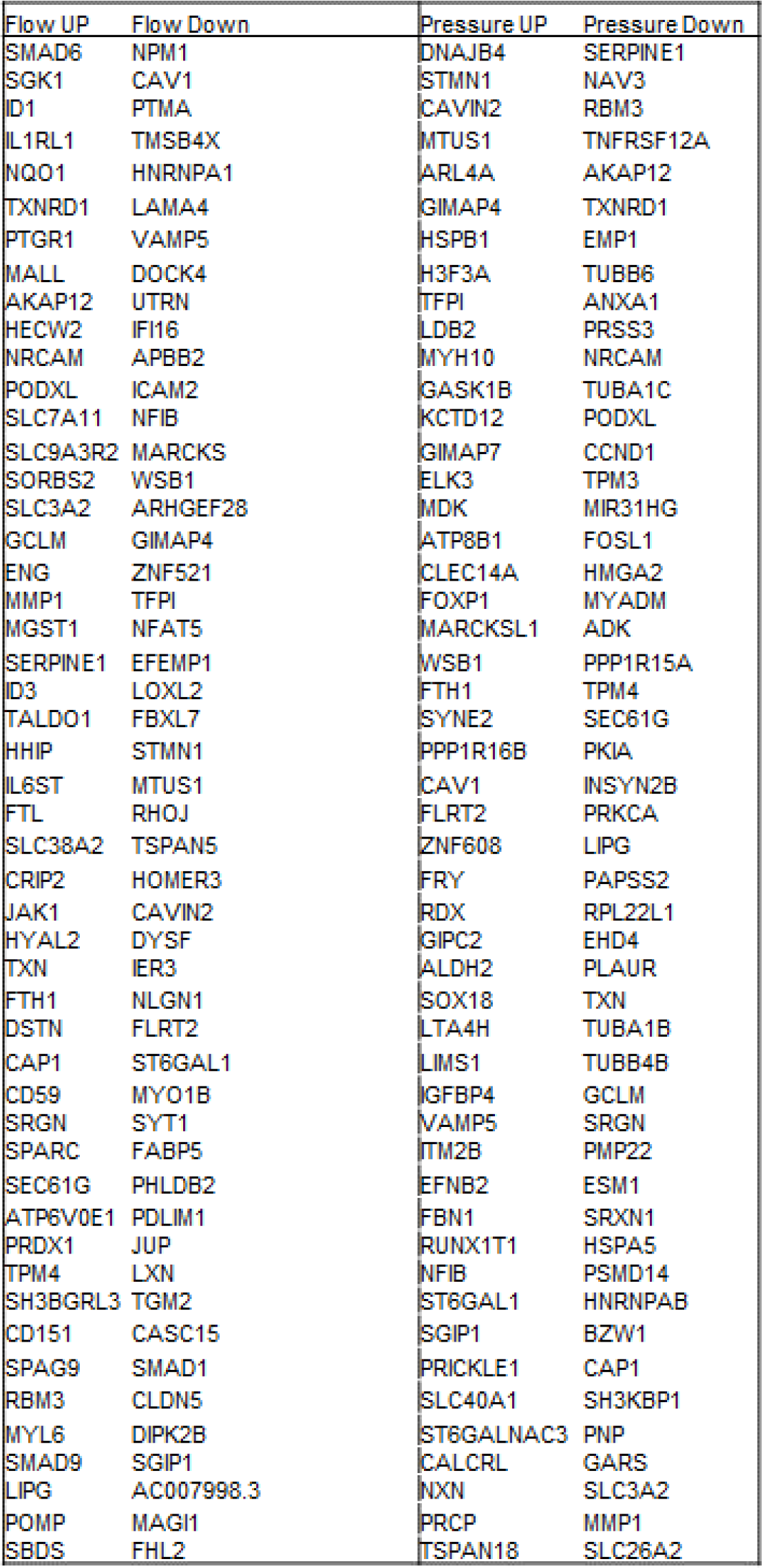
Top 50 differentially expressed genes regulated by flow and pressure in scRNAseq analysis.

Next, focusing on samples under constant flow but varying pressure (restricted to non-proliferating cells of cluster 1), we evaluated the transcriptional response to pressure (Fig. 2C). Differential gene expression analysis between ECs exposed to the highest (P=60 mmHg) vs lowest (P=0 mmHg) pressure conditions unveiled a distinct gene-set signature indicative of pressure response in single-cell data (Fig. 2C, Table 1). A significant dose-response effect of gene-set scores for pressure-responsive genes was evident across all pressure conditions (P=0, 20, and 60-mm Hg). Given the known involvement of the MYC pathway negatively regulating arterialization ^39,40^ and endothelial cell quiescence ^41,42^, we investigated hallmark MYC target genes under varying pressures. Genes within the MYC pathway exhibited dose-dependent downregulation in response to pressure, suggesting that elevated pressure levels may alter the metabolic and proliferation states of ECs through MYC.

A conserved pressure-dependent gene expression program operates during arterial and hemogenic EC maturation in the mouse aorta.

Endothelial cells in the developing embryonic aorta experience dynamic shifts in hemodynamic forces, including flow/shear stress and hydrostatic pressure, initiated by the onset of the heartbeat ^7,43^. In our recent publications ^44,45^, we presented single-cell transcriptional analysis of EC maturation towards arterial and hemogenic fates in the murine aorta, spanning the developmental period from embryonic day 8.5 (E8.5) to E11.5. This in vivo murine aorta scRNA-seq data reveals a correlation between flow and pressure-dependent transcriptional programs and gene-expression signatures specific to arterial ECs and Hematopoietic Stem Cells (HSCs). Notably, genes identified as differentially expressed in HUVECs under high pressure (60 mmHg) versus low pressure (0 mmHg) conditions in vitro (Fig. 2C, Table 1) were significantly upregulated in late stage (E10.5 to E11.5, high pressure) arterial/hemogenic EC compared to the early stage (E8.5-E9.5, low pressure), and also in ECs transcriptionally identified as arterial (high pressure state) verses venous (low pressure state) in the mouse aorta scRNA-seq data (Fig. 2D). The expression changes in pressure-responsive gene sets further correlated with downregulation of MYC target genes previously implicated in arterial EC maturation ^39^ and associated with maturation of hemogenic ECs with hematopoietic stem cell fate in the embryonic aorta ^45^ (Fig. 2D). These findings suggest a conserved pressure-responsive gene signature unveiled by our in vitro HUVEC microvessel models that may be activated during normal development in the nascent aorta, contributing to arterial and hemogenic EC maturation.

## Discussion

Biomechanical cues emerge as pivotal regulators during embryonic development, particularly in the cardiovascular system. From the initiation of blood flow, hemodynamic forces such as wall shear stress, cyclic strains, and hydrostatic pressure exert influence on vascular wall cells, in particular the endothelium, shaping their cell fate, structure and function.^7,46^ ECs, as the dynamic interface between the blood and the local tissue, possess the ability to rapidly sense alterations in these hemodynamic forces, and engage in signaling and secretion, thereby contributing to hemodynamic regulation.^47^ While previous studies have predominantly examined EC response to shear flow irrespective of hydrostatic pressure, the combined effects of pressure and shear forces remain less understood. In this study, we developed a platform to expose ECs to controlled, combinatorial shear and pressure conditions, revealing that pressure modulates both morphologic and transcriptional responses under varying shear stress conditions. This observation underscores the significant role of non-shear forces in endothelial biology and suggests that an integrated perspective on mechanotransduction could provide new insights into EC behavior.

We noted that pressure conditions substantially influenced the shape and orientation of cells in response to flow (at shear stress of 5 dyne/cm^2^). Under low pressure, ECs exhibited typical responses, elongating and aligning in the flow direction with notable clustering of Notch1-ECD at the downstream pole. Conversely, at higher pressures, ECs displayed a reversal of these morphologic trends, appearing smaller and oriented perpendicular to the flow direction. This suggests that pressure signaling may supersede shear-sensing pathways under certain conditions. Furthermore, our findings indicated that high pressure led to a higher cell density of ECs, while a similar fraction of cells showed positive for Ki67 under low pressure, suggesting a potential initial burst in proliferation followed by contact inhibition at high pressure.

Recent studies have shown that hydrostatic pressure independent of flow can transiently activate the Ras/extracellular signal–regulated kinase (ERK) and protein kinase C (PKC) pathways in endothelial cells, promoting tubulogenesis.^48^ Here we demonstrated that hydrostatic pressure altered the transcriptional profiles of ECs in response to flow via both bulk and single cell RNA sequencing. While fluid shear stress predominantly drove changes in EC transcriptional profiles, consistent with prior studies,^7,19^ we showed that pressure conditions modified the EC response to flow. High-pressure appeared to downregulate genes associated with GTPase signaling and cell morphogenesis, while upregulating genes associated with cell proliferation, suggesting opposing effects of pressure and flow on genes involved in mechanosensory signaling and cytoskeletal organization. In our scRNAseq analysis, we also identified a distinct set of genes that exhibited a dose-dependent response to pressure in ECs under fixed flow shear. Particularly, genes involved in the MYC pathway, which regulates endothelial metabolism, arterial and hemogenic maturation, and the establishment and maintenance of functional vascular structures,^49–53^ were downregulated as hydrostatic pressure increased under constant flow. Thus, our results provide evidence supporting a role for hydrostatic pressure in modulating EC identity and function, potentially through the modulation of the MYC pathway.

Interestingly, when we examined scRNAseq data from murine embryonic ECs isolated during a developmental window encompassing the onset of circulation,^44,45^ we found that pressure-responsive gene sets defined experimentally in our engineered vessels were upregulated as a function of developmental stage, correlating with expected changes in pressure. Similar changes in pressure-responsive genes were observed when comparing transcriptionally defined venous EC (under low pressure conditions) versus arterial/hemogenic EC (under high pressure conditions). Consistent with our observations in the engineered microvessel conditions, these transcriptional changes in pressure-responsive genes correlated with downregulation of MYC target genes implicated in arterial and hemogenic EC maturation in the murine embryo.^39,45^ These findings suggest a conserved, pressure-responsive gene signature activated during normal development, potentially contributing to arterial and hemogenic EC maturation.

Together, these studies highlight the developmental significance of an integrated endothelial response to both flow and pressure-dependent transcriptional programs, suggesting an important role in promoting distinct endothelial cell fates. Our bulk and scRNAseq data also provide a valuable resource for future studies aimed at elucidating the mechanisms by which ECs sense and integrate flow shear stress and pressure to activate downstream signaling pathways governing their fates.

While our studies represent a significant advance in understanding the combinatorial response of ECs to pressure and flow, they have several notable limitations. Firstly, the absence of physiological stretch in our experimental setup is a constraint. Previous studies have demonstrated that ECs subjected to physiological strain align perpendicular to the strain direction, reinforcing shear-induced alignment in the flow direction.^54,55^ In vivo, pressure and stretch are intricately linked, and their combined effect on EC behavior is complex. The absence of this factor in our system may have influenced our observations, particularly in terms of cell alignment and morphological responses. Another limitation is the lack of substrate remodeling with the use of polydimethylsiloxane (PDMS) for tube fabrication and high-pressure exposure.

Future utilization of cell-remodeling matrices with different mechanical properties and cellular compositions on the vessel wall could yield richer insights into multifactorial effects on endothelial cell response to flow. Despite these limitations, our study paves the way for further research into the integrated response of endothelial cells to hemodynamic forces, with implications for understanding vascular development and disease progression.

## Acknowledgement

We acknowledge the Lynn and Mike Garvey Microscopy Imaging Core in the Institute for Stem Cell and Regenerative Medicine. We thank Taylor Merkel’s assistance in collecting part of the data. This work was supported by funding from the NIH 1R61/33HL154250, R01AI148802 and UG3 TR003288 (YZ), R01HL168110 (BH) and UW ISCRM IPA award (YZ and BH). T.I is supported for overseas medical research by Takeda Science Foundation, The Nakatomi Foundation and Kitasato University School of Medicine Alumni Association. We also acknowledge the Washington Nanofabrication Facility / Molecular Analysis Facility, a National Nanotechnology Coordinated Infrastructure (NNCI) site at the University of Washington with partial support from the National Science Foundation via awards NNCI-2025489 and NNCI-1542101.

## Author Contribution

C.M., B.H. and Y.Z. conceived and designed the study. C.M. and T.I. carried out the experiments with the assistance of S.R. and A.H. C.M. carried out bioinformatic analysis for bulk RNAseq, whereas T.I. and B.H. carried out relevant bioinformatic analysis for scRNAseq. All authors contributed to writing and editing the paper.

## Methods

### Microfluidic channel design and fabrication for decoupling the pressure and flow exposure

To manipulate pressure and flow conditions, we developed a perfusion setup with multiple microfluidic channels, with the primary channel for cell culture whereas downstream channels as resistors to elevate pressure in the culture channel. For cell culture, we fabricated a single channel (w = 500 μm, h = 100 μm, L = 20 mm) with higher aspect ratio was designed to minimize edge effects for brightfield and immunofluorescence images of ECs. In cultures for sequencing analysis, we fabricated a single tube (230 μm diameter, 20 mm length) using needle subtraction in PDMS, to eliminate edge effects and ensure uniform stress distribution, and accommodate approximately 5000 cells. Resistance channels (100 μm width, 100 μm height) were designed to provide resistance corresponding to pressures of 20 mmHg or 60 mmHg at flow rates yielding shear stresses of 5 and 10 dyne/cm^2^ in the culture channels. Shear stress for circular tubes was estimated by equation [inline], where Q is the volumetric flow rate, µ is the viscosity, and r is the channel radius. The pressure drop in all channels was estimated by equation [inline], where L is the channel length.

Microfluidic resistor channels were designed using LayoutEditor (juspertor GmbH) and fabricated in silicon using standard lithography. Briefly, chrome masks of channel designs were exposed on a DWL 66+ (Hiedelberg GmbH) and etched. Silicon wafers were spin coated with AZ9260 (Microchem, thickness ≈ 6 μm), exposed using a contact aligner and chrome masks (ABM), and developed with AZ400K (Microchem). Patterned wafers underwent deep reactive ion etching (SPTS Rapier DRIE), followed by resist stripping (EKC, Dupont). Final channel dimensions were measured optically (Keyence VK-X150K) and physically (KLA Tencor P15 Contact Profilometer).

To transfer the finished patterns to PDMS, silicon wafers were silanized with trichloro(3,3,3- trifluoropropyl) silane (Sigma). A 10:1 mixture of PDMS base and curing agent (Sylgard 184, Dow) was poured over the wafer and cured at 65 °C for at least 3 hours. Individual channels were separated, and inlet and outlet holes were punched with a 2 mm biopsy punch (Integra Miltex). Patterned PDMS channels were bonded to cover glass (Thorlabs #1.5) or plain glass slides (Fisher Scientific) using air plasma (Power = 75 W, Plasma Etch) and annealed at 65 °C for at least 1 hour.

To validate the hemodynamic properties of fabricated channels we constructed a flow circuit driven by a syringe pump (KD Scientific). Syringes were connected to a microfluidic flow sensor (Elveflow MFS-80), a bare PDMS culture tube, a microfluidic pressure sensor (Elveflow MPS-1), and bare PDMS resistors of different lengths. Perfusion media (EGM2 + 3.5% dextran) was loaded into syringes and run through the circuit. Steady-state values for flow rate and pressure were recorded using an Elveflow controller (Elveflow OBS).

### Channel seeding and culture

Prior to cell culture, channels were sterilized by 30 minutes of UV exposure and coated with human fibronectin (5 μg/mL) for 1 hour at 37 °C. Channels were rinsed with fresh culture medium (EGM2, Lonza) and seeded with human pulmonary artery endothelial cells in rectangular channel experiments (HPAEC, Lonza, Fig.1A-D), or human umbilical vein endothelial cells in round channel experiments (HUVEC, Lonza, Fig 1E-H) by perfusion of 10 μL of a 4 million cells/mL suspension. Cells adhered for 1 hour at 37 °C before rinsing with fresh media to remove unattached cells. Attached cells in inlet and outlet reservoir were removed by scraping these regions with a gel loading pipette tip attached to a vacuum aspirator, leaving cells only in the channel cross-section. Channels equilibrated overnight in an incubator with gravity driven media culture through the inlet and outlet.

### Channel perfusion

The perfusion medium, consisting of normal culture medium supplemented with 3.5% dextran to mimic the viscosity of blood (∼3.5 cP), was degassed under vacuum for 30 minutes, connected to culture channels. Flow rates were adjusted to achieve the desired shear stress and channels were perfused for 24 hours. For the selected tube, flow rate is set as 9.5 µL/min to achieve shear stress of 5 dyne/cm^2^, Length of resistor channel is 10.7mm and 34.5mm to achieve outlet pressure of 20 and 60 mmHg respectively.

### Immunofluorescent staining and imaging

Culture channels were disconnected from the syringe pump and immediately perfused with 250 μL of 4% paraformaldehyde for 10 minutes, followed by washing with PBS (3 x 5 min). For staining, channels were blocked and permeabilized with 2% BSA and 0.5% Triton X-100 in PBS for 30 minutes. Channels were then incubated with primary antibodies for 1 hour at room temperature or overnight at 4 °C, followed by PBS washes. Secondary antibodies, along with actin and nuclear stains, were added for 1 hour at room temperature before washing with PBS. The primary antibodies used included: VECAD/CD144 (1:100, Abcam ab33168), AQP-1 (1:100, Santa Cruz sc-25287), GJA4 (1:100, Abcam ab181701), DLL4 (1:100, Novus NB600-892), Ki67 (1:100, Abcam ab16667), Notch1-ICD (1:100, Cell Signaling 3608S). Secondary antibodies and other stains included: Notch1- ECD (1:100, BD 566023), DLL1 (1:50, Biolegend 346403), VECAD/CD144 (1:100, eBioscience 17-1449-42), Alexa Fluor goat anti-rabbit 647 (1:100, Thermo Fisher A-21235), Alexa Fluor goat anti-mouse 568 (1:100, Thermo Fisher A-11004), Alexa Fluor Phalloidin 488, 568, and 647 (1:100, Thermo Fisher Scientific, 488:A12379, 568:A12380, 647:A22287), and Hoescht 33342. Stained channels were imaged with a widefield microscope (Nikon Eclipse Ti2) and a line scanning confocal system (Nikon A1R).

### Image analysis, quantification and statistics

Images were processed using Fiji.^45^ For cell morphology analysis, maximum intensity projections of Z-stacks of cells adhered to the channel luminal walls were used to draw cell borders from junctional or actin stains. Cell size, perimeter, angle, and shape descriptors were calculated and exported as csv files for analysis in R. Cell orientation was determined using the fit ellipse function to identify its major and minor axis with the cell angle defined as the angle between the flow direction (0 degrees) and the cell’s major axis. At least 50 cells identified across the 3 fields were analyzed for each channel. Nuclear counts were obtained by thresholding the Hoescht signal and utilizing the “Analyze Particles” function in Fiji. Ki67 positive nuclei were manually counted to determine the percentage of Ki67 positive nuclei. Individual cell parameters were compared based on pressure condition using an unpaired two-sided Wilcoxon Rank Sum test (significance set at p < 0.05). Nuclei counts and the ratio of Ki67 positive nuclei were averaged for each channel replicate (n = 3 channels) and compared using an unpaired two-sided Student’s t-test (significance set at p < 0.05).

### RNA isolation, sequencing, and analysis

After 24 hours of flow, cultures of endothelial cells were disconnected from the syringe pump, and 350 μL of lysis buffer (RLT + 1% β- mercaptoethanol, Qiagen) was perfused through each channel. RNA was isolated using a RNeasy Micro kit (Qiagen) with DNase treatment. Isolated RNA was then submitted to the Fred Hutchinson Genomics Resource for quality control, library preparation, and sequencing. RNA quality was confirmed (RINe ≥ 9.7) using an Agilent Tapestation. Libraries were prepared using SMART-Seq v4 Ultra Low Input (Takara) and Nextera XT DNA Library Preparation (Illumina) kits, and sequenced on an Illumina NovaSeq 6000 (paired end, 50 bp). Reads were aligned to hg38 using STAR ^56^ and counts were generated using the featureCounts function of the subRead package.^57^ Quality control was performed using RSeQC^58^ and differential expression analysis was performed using edgeR (3.28.1).^59^ Genes were considered differentially expressed if the P and FDR values were ≤ 0.05 and the absolute log fold change was ≥ 0.585. Gene ontology and gene set enrichment analysis were conducted using clusterProfiler^60^, and the Molecular Signatures Database^61^. All data has been deposited at NCBI GEO (GSE260600).

### Single cell RNA sequencing and analysis

HUVECs cultured for 24 hours under constant flow and varying pressure conditions were harvested by trypsinization from multiple devices for each condition. Approximately 50,000 cells from each culture condition were pooled, resuspended in 0.04% ultrapure BSA (Invitrogen) in PBS on ice for scRNAseq. Cells from each sample were labeled with TotalSeq™-B0251-0254 anti-human Hashtag Antibodies (clone LNH-94, Biolegend, RRID: AB_ 2814347-2814350, at a concentration of 125 mg/ml) for multiplexing^62^ following the manufacturer’s protocol (10X Genomics, Cell Surface Protein Labeling for Single Cell RNA Sequencing Protocols with Feature Barcoding technology). Hashtagged cells were then pooled and loaded onto the Chromium Single Cell Chip G and processed through the controller (10X Genomics). Single cell mRNA libraries were generated with the 10X Chromium Next GEM Single Cell3’ Reagent Kits version 3.1 with Feature Barcoding technology for Cell Surface Protein, according to 10X protocol. Sequencing was performed for pooled libraries on an Illumina NextSeq 2000 using the 100 cycle, high output kit, targeting a minimum of 40,000 reads per cell. Raw and processed sc-RNA-seq data have been deposited at NCBI GEO (Accession number GSE261063).

### Single cell RNA sequencing analysis and quality control

The Cell Ranger 7.1.0 pipeline (10X Genomics) was used for alignment, demultiplexing, and generating the feature barcode matrix. Monocle3 (v.3.1.2.9) ^63,64^ was used for downstream analysis, combining normalized data from each sample. Cells with high (>10%) mitochondrial reads, low genes counts (<1,000) or low UMI counts (UMI<5,000) were excluded. Uniform Manifold Approximation (UMAP) was used for dimensionality reduction^65^. The alignCDS() function was applied to remove batch effects ^66^. Cells were clustered by the Louvain method implemented in Monocle3 ^67^.

Differential gene expression analysis used regression models with the fit_models() and coefficient_table() functions in Monocle3, to identify genes that were differentially expressed between experimental conditions based on significance values (q value) adjusted for multiple hypothesis testing using the Benjamini and Hochberg correction method (full gene lists with q-values provided in Table 2). Gene set scores were calculated for each single cell as the log-transformed sum of the size factor-normalized expression for each gene in canonical/published signature gene sets, experimentally determined genes sets (flow and pressure-dependent), or from the Molecular Signatures Database (https://www.gsea-msigdb.org/gsea/msigdb/index.jsp) (see Table 2 for gene sets used in this study). Violin plots of gene set scores based on sample/condition, embryonic stage, or cell type were generated using the ggplot functions geom_violin and geom_boxplot (boxplots show median values and interquartile ranges; upper/lower whiskers show 1.5X interquartile range).

**Table 2:**
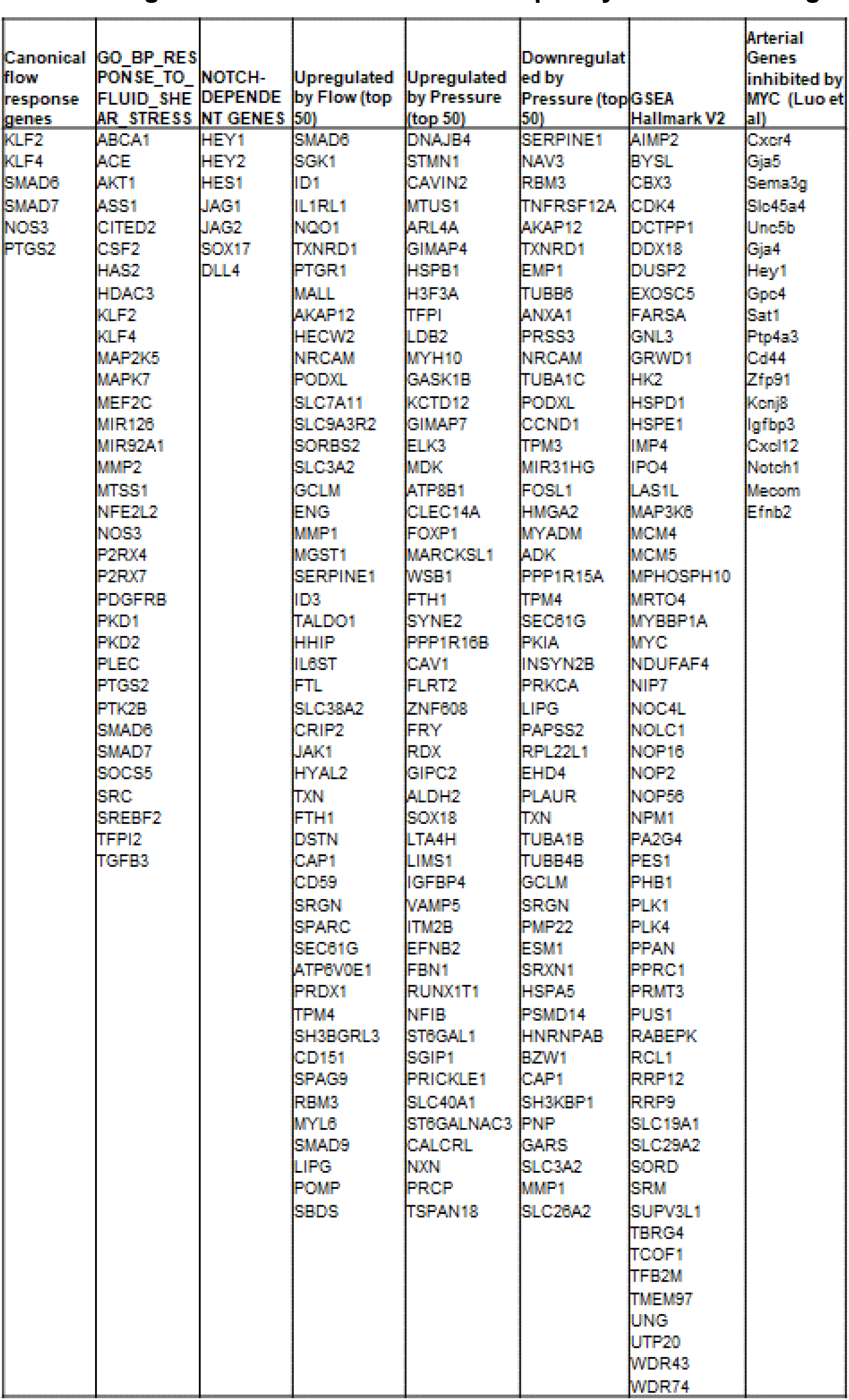
Gene list and gene sets defined in sc RNAseq analysis shown in. **Figure 2**.

Statistical analysis used the Wilcoxon Rank Sum Test (ggupbr package v0.4.0) to calculate p values as indicated. Our previously published scRNAseq data from EC and hematopoietic populations isolated from early mouse embryos encompassing embryonic stages E8.5 to E11.5 ^44,45^ were combined in a single cell data set for downstream analysis in Monocle3. Cell type classification was performed using the Garnett package (v.0.2.15) in Monocle 3. Marker gene sets based on established cell type-specific genes (listed below) were used to train a classifier data (using the train_cell_classifier function with default settings) and classify cell types (using the classify_cells function). Cells not classified as EC/HE were excluded for analysis in this study.

| Cell Type | Genes Expressed | Genes not expressed |
| --- | --- | --- |
| Venous EC | *Nr2f2, Nrp2* | *Hey2, Gja5, Myb, Spn* |
| Arterial EC/HE | *Gja5, Hey2* | *Nr2f2, Nrp2, Myb, Spn* |
| Hematopoietic | *Myb, Spn* | *Hey2* |

R scripts used for sc-RNAseq analysis are available at the Fred Hutchinson Cancer Center GitHub: https://github.com/FredHutch/Mandrycky-etal-2024

## Supplementary Figures

**Supplementary Figure 1.**
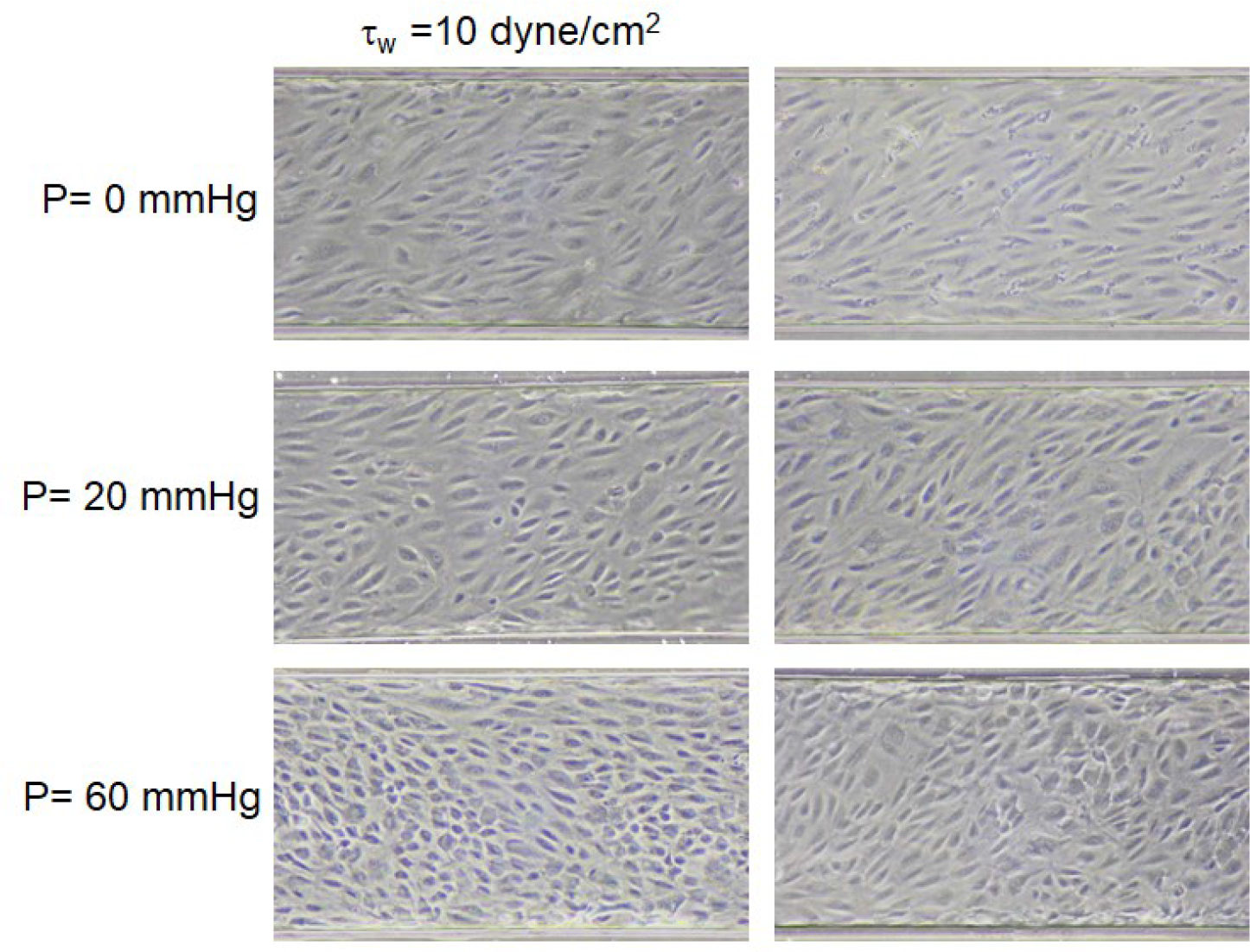
Brightfield images of HPAECs cultured under flow shear (wall shear stress at 10 dyne/cm^2^ and pressure at 0, 20, and 60mmHg.

**Supplementary Figure 2.**
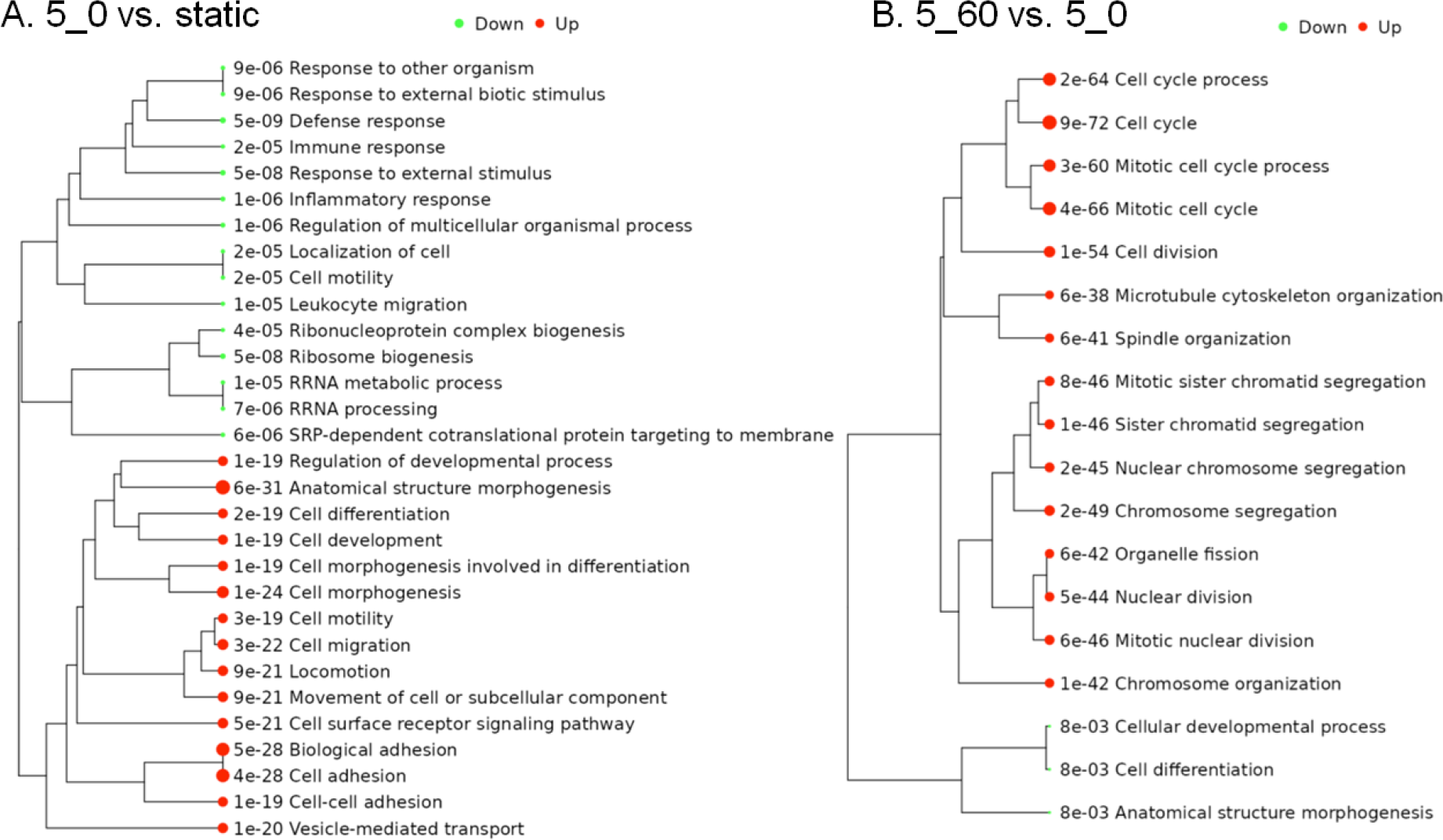
Pathway trees for (A) flow driven effect comparing shear stress at 5 dyne/cm^2^ vs. static condition, and (B) pressure driven effect comparing 60mmHg vs. 0 under flow shear of 5 dyne/cm^2^.

**Supplementary Fig. 3.**
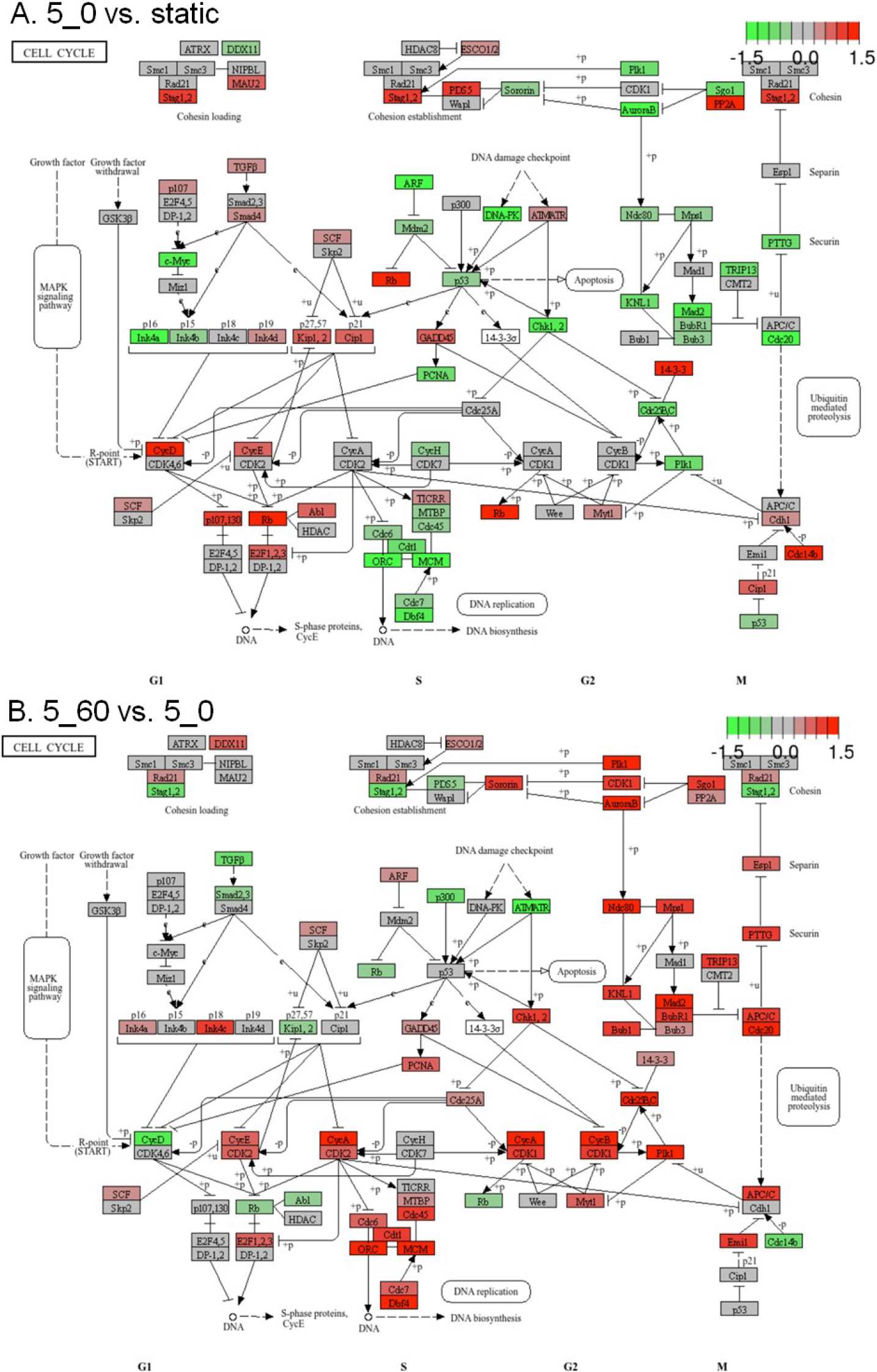
Cell cycle gene network comparison for (A) shear stress at 5 dyne/cm^2^ vs. static condition, pressure at nearly zero, and (B) pressure at 60mmHg vs. 0 under flow shear of 5 dyne/cm^2^.

